# A comprehensive genomics solution for HIV surveillance and clinical monitoring in a global health setting

**DOI:** 10.1101/397083

**Authors:** David Bonsall, Tanya Golubchik, Mariateresa de Cesare, Mohammed Limbada, Barry Kosloff, George MacIntyre-Cockett, Matthew Hall, Chris Wymant, M Azim Ansari, Lucie Abeler-Dörner, Ab Schaap, Anthony Brown, Eleanor Barnes, Estelle Piwowar-Manning, Ethan Wilson, Lynda Emel, Richard Hayes, Sarah Fidler, Helen Ayles, Rory Bowden, Christophe Fraser

## Abstract

High-throughput viral genetic sequencing is needed to monitor the spread of drug resistance, direct optimal antiretroviral regimes, and to identify transmission dynamics in generalised HIV epidemics. Public health efforts to sequence HIV genomes at scale face three major technical challenges: (i) minimising assay cost and protocol complexity, (ii) maximising sensitivity, and (iii) recovering accurate and unbiased sequences of both the genome consensus and the within-host viral diversity. Here we present a novel, high-throughput, virus-enriched sequencing method and computational pipeline tailored specifically to HIV (veSEQ-HIV), which addresses all three technical challenges, and can be used directly on leftover blood drawn for routine CD4 testing. We demonstrate its performance on 1,620 plasma samples collected from consenting individuals attending 10 large urban clinics in Zambia, partners of HPTN 071 (PopART). We show that veSEQ-HIV consistently recovers complete HIV genomes from the majority of samples of different subtypes, and is also quantitative: the number of HIV reads per sample obtained by veSEQ-HIV estimates viral load without the need for additional testing. Both quantitativity and sensitivity were assessed on a subset of 126 samples with clinically measured viral loads, and with standardized quantification controls (VL 100 – 5,000,000 RNA copies/ml). Complete HIV genomes were recovered from 93% (85/91) of samples when viral load was over 1,000 copies per ml. The quantitative nature of the assay implies that variant frequencies estimated with veSEQ-HIV are representative of true variant frequencies in the sample. Detection of minority variants can be exploited for epidemiological analysis of transmission and drug resistance, and we show how the information contained in individual reads of a veSEQ-HIV sample can be used to detect linkage between multiple mutations associated with resistance to antiretroviral therapy. Less than 2% of reads obtained by veSEQ-HIV were identified as *in silico* contamination events using updates to the *phyloscanner* software (*phyloscanner clean*) that we show to be 95% sensitive and 99% specific at ‘decontaminating’ NGS data. The cost of the assay — approximately 45 USD per sample — compares favourably with existing VL and HIV genotyping tests, and provides the additional value of viral load quantification and inference of drug resistance with a single test. veSEQ-HIV is well suited to large public health efforts and is being applied to all ∼9000 samples collected for the HPTN 071-2 (PopART Phylogenetics) study.

## 1. Introduction

Achieving sustained reductions in the incidence of HIV infections through programmes of universal access to testing and antiretroviral treatment (UTT) remains a major goal in public health. Current international focus has been on working towards the UNAIDS ‘90-90-90’ targets: 90% of people living with HIV (PLWH) diagnosed, 90% of those on antiretroviral therapy, and 90% of those successfully virally suppressed [1]. However, this leaves 27% of PLWH not virally suppressed and still at risk of transmitting. There is substantial heterogeneity in risks of both becoming infected and transmitting HIV; if certain traits that associate with HIV transmission (such as high sexual partner exchange rates, and low rates of accessing sexual health services) are disproportionately found in the virally unsuppressed population, this will undermine the ability of the 90-90-90 goals to control the epidemic [2] [3]. Consistent with this concern, an interim analysis of the HPTN 071 (PopART) cluster-randomized trial of UTT in Zambia found the percentage of individuals who knew their HIV+ status was lowest in men and in younger people [34]. There is therefore urgent need to study transmission patterns and identify any correlates of enhanced transmission risk that could be specifically targeted for intervention. The success of UTT programmes is also threatened by drug resistance, which is expected to increase in frequency as treatment is scaled up [4]. A 2017 report by the WHO identified parts of the world where more than 10% of people living with HIV already harbour virus resistant to antiretroviral compounds [5].

Both transmission patterns and the spread of drug resistance can be better understood using viral sequence data. To date, clinical drug resistance testing has primarily relied on Sanger consensus sequencing of *pol* genes. In high-income countries, where drug resistance testing is routinely performed to guide choice of effective ART, *pol* sequences have been used to identify transmission clusters [6]. The potential for viral whole-genome sequencing to transform global health surveillance operations has been noted [7] and implementation would benefit from the higher throughput of next-generation sequencing (NGS) technologies. NGS also produces detailed minority variant information, which can enhance resolution in transmission analyses, indicate direction of transmission [8], and detect low-frequency drug-resistant viral variants. Despite its benefits, adoption of NGS for HIV has been slow, in part due to technical difficulties in obtaining whole genome sequences for all genotypes particularly at low viral loads, and uncertainty over distinguishing low-frequency mutations from the sequencing artefacts and contamination that occur during massively-parallel sequencing. And although costs are lower than Sanger consensus sequencing, they remain prohibitively high for wide-scale use in low and middle-income countries.

Large-scale sequencing of HIV genomes with NGS using overlapping 2-3 kb PCR amplicons [9] has been used to produce complete genomes for European samples [10], but the method’s performance was found to be far from optimal on sub-Saharan African samples, with amplification failures resulting in biased genome coverage [11]. In our previously reported protocol, veSEQ, for Hepatitis C Virus (HCV), a virus which is even more diverse than HIV, we avoided such a genotype-dependent failure rate by avoiding virus-specific PCR altogether [12]. Oligonucleotide probes of 120 bp were designed to capture the known epidemic diversity of HCV; we found that these probes could effectively capture target material with at least 80% nucleotide homology, in a manner that was unbiased with respect to diversity within this range (0 to 20%). The probes were used to enrich viral sequences from total RNA-seq libraries generated from plasma. We have since translated this principle to other viruses (Zika virus [13] ; HBV, in progress).

Here we describe our adoption of this sequencing method for HIV, optimising both genome coverage and average fragment size (the latter enhancing the resolution of within-host phylogenetics). We report our method, veSEQ-HIV, a comprehensive laboratory and computational pipeline, able to distinguish true minor variants, including low-frequency drug resistance mutations, from sequencing artefacts. We show our method to be quantitative – capable of estimating viral load across at least five orders of magnitude – and obtain whole genome sequences at viral loads as low as 100 RNA copies per ml. We describe our optimisations to minimise the bias inherent in PCR amplification, while also minimising quantities of reagents, reducing their cost to one fifth of the WHO 2015 budget recommendation for both viral load measurements and drug resistance testing [14].

We demonstrate the method’s performance using the first 1,620 samples collected from consenting invividuals attending a government ART clinic within 9 of the HPTN 071 (PopART) trial communities. The PopART phylogenetics study (HPTN-071-2) is an ancillary study to the main PopART clinical trial; the largest ever cluster-randomized trial of HIV prevention that compares standard of care to a comprehensive intervention package of proactive home-based testing, rapid linkage to care, immediate initiation of ART and additional standard preventative measures [15].

## 2. Methods

### 2.1. Patient population

Patients were recruited to the HPTN 071-2 (PopART phylogenetics) study by research assistants at ten urban primary healthcare facilities, located in nine of the twelve Zambian communities of the main trial (one community had two health care facilities) [15]. The nine communities involved were evenly split between the three study arms of HPTN-071. Patients were recruited if they were aged 18 or over, if not currently taking ART, and if they specifically consented to the ancillary phylogenetic study. Most patients were either newly enrolled in the clinic, or enrolled and newly eligible for ART; a small fraction were recruited having recently missed several doses of ART.

The study protocol is available here [https://www.hptn.org/sites/default/files/inlinefiles/HPTN%20071-2%2C%20Version%202.0%20%2807-14-2017%29.pdf], and has been approved by the ethics committees of the University of Zambia (c/o the Zambian ministry of health) and of the London School of Hygiene and Tropical Medicine.

### 2.2. Sampling

No additional blood sample was required for this ancillary study. Saved unused samples of blood collected from consenting individuals undergoing routine CD4 testing were transported to the local hospital, usually on the same day. Blood was centrifuged twice and two 500 *µ*L aliquots of plasma were frozen at -80 °C. Samples were transported to a central research laboratory (ZAMBART facility) in Lusaka, Zambia, using a mobile -80° C freezer, and then shipped to the sequencing laboratory in the UK. Samples were processed approximately in order of collection, and represent the diversity of the population recruited at the beginning of the study (Figure 2).

**Figure 1:**
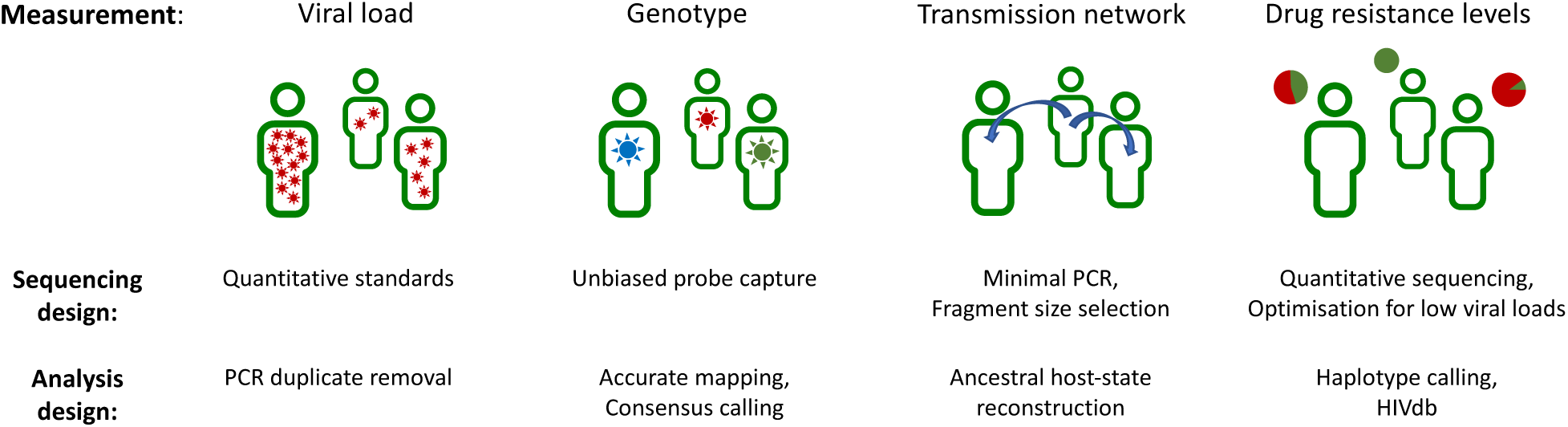
Information included in theveSEQ-HIV method. TheveSEQ-HIV method was developed to provide multiple measurements from a single assay, including viral load, genotype, inference of the transmission events and drug resistance levels.

**Figure 2:**
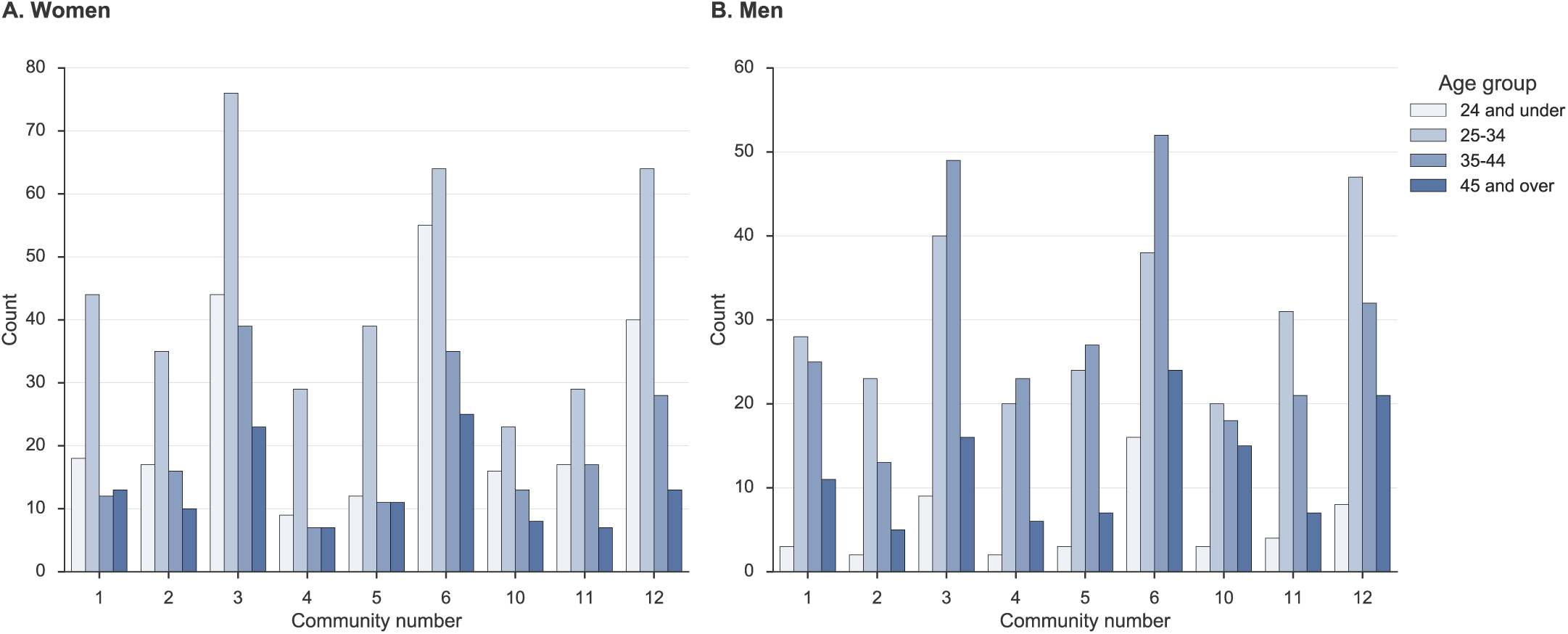
Study participants with sequenced HIV samples, by HPTN-071 (PopART) community number. Panel A, Women; Panel B, Men. Sequenced samples came from the nine communities in the 071-2 ancillary study in Zambia, with differences in total numbersreflecting differences in size of community catchment area and number of participating healthcare facilities. All three study arms were equally represented among the nine communities.

### 2.3. Laboratory methods

Total RNA was extracted with magnetised silica from HIV-infected plasma lysed with guanidine thiocyanate and with ethanol washes and elution steps performed using the NUCLISENS easyMAG system (bioMérieux). The total 30 *µ*l elution was reduced in volume with Agencourt RNAClean XP (Beckman Coulter) to maximize the input RNA mass while minimizing volume for library preparation.

Libraries retaining directionality were prepared using the SMARTer Stranded Total RNA-Seq Kit v2 — Pico Input Mammalian (Clontech, Takara Bio) with the following protocol specifications. Total RNA was first denatured at 72°C with the addition of tagged random hexamers to prime reverse transcription, followed by cDNA synthesis according to the manufacturer protocol option with no fragmentation. The first strand cDNA was then converted into double-stranded dual-indexed amplified cDNA libraries using in-house sets of 96 i7 and 96 i5 indexed primers [16], using a maximum of 12 PCR cycles. All reaction volumes were reduced to one quarter of the SMARTer kit recommendation and set up was either prepared manually, or automated using the Echo 525 (Labcyte) low-volume liquid handler.

No depletion of ribosomal cDNAwas carried out prior to target enrichment. Equal volumes (5 *µ*l from a total of 12.5 *µ*l) of each amplified library were pooled in 96-plex without prior clean-up. The pool was cleaned with a lower ratio of Agencourt AMPure XP than recommended by the SMARTer protocol, to eliminate shorter libraries (0.68X). The size distribution and concentration of the 96-plex was assessed using a High Sensitivity D1000 ScreenTape assay on a TapeStation system (Agilent) and a Qubit dsDNA HS Assay (Thermo Fisher Scientific).

A total of 500 ng of pooled libraries was hybridized (SeqCap EZ Reagent kit, Roche) to a mixture of custom HIV-specific biotinylated 120-mer oligonucleotides (xGen Lockdown Probes, Integrated DNA Technologies), ten pulled down with streptavidin-conjugated beads as previously reported [12]. Unbound DNA was washed off the beads (SeqCap EZ Hybridization and Wash kit, Roche), and the captured libraries were then PCR amplified to produce the final pool for sequencing using a MiSeq (Illumina) instrument with v3 chemistry for a read length up to 300 nt paired-end. Alternatively, upto 384 samples were sequenced on HiSeq 2500 set to Rapid run mode using HiSeq Rapid SBS Kit v2 with maximum read lengths of 250 nt.

### 2.4. Clinical viral loads

Clinical viral load measurements were obtained within the Oxford University Hospital’s clinical microbiology laboratory using the COBAS^®^ AmpliPrep/COBAS^®^ TaqMan^®^ HIV1 Test (Roche Molecular Systems, Branchburg, NJ, USA).

### 2.5. Analytics

Raw sequencing reads were first processed with Kraken [17] to identify human and bacterial reads. Kraken was run with default parameters (k=31 with no filtering), using a custom database containing the human genome together with all bacterial, archaeal and viral genomes from RefSeq, a subset of fungal genomes, and all 9,049 complete HIV genomes from GenBank (last updated 18 May 2018). Reads were filtered to retain only viral and unclassified sequences, and these were trimmed to remove adaptors and low-quality bases using Trimmomatic [18], retaining reads of at least 80 bp. Filtered, trimmed sequences were assembled into contigs using SPAdes [19] and metaSPAdes [20] with default parameters for k (21 to 127). Contiguous sequences assembled from both assembly runs were clustered using cd-hit-est to remove redundant contigs [21], retaining the longest sequence in each cluster with minimum sequence identity threshold of 0.9. Contigs together with the filtered reads were then used to generate HIV genomes and variant frequencies using shiver [22], with position-based deduplication of reads enabled. Samples for which no contigs could be assembled were mapped to the closest known HIV reference as identified by Kallisto [23], hashing the filtered reads against a set of HIV references downloaded from the Los Alamos HIV database (http://www.hiv.lanl.gov/), and taking the closest matching genome as the mapping reference for shiver. QC metrics from each processing step are presented in Supplementary Table 1.

Sequence-derived viral load, in copies per ml, was calculated from the number of deduplicated HIV reads for each sample, using the regression slope and intercept determined from the clinically measured viral load in a subset of 126 samples. To ensure a defined value of the log transformation for samples with zero reads, a pseudocount of 1 was added to the read number. The equation was:

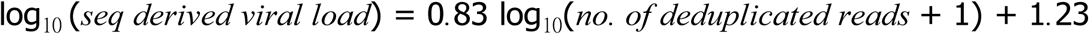

Contaminant reads were identified and removed using *phyloscanner* for in-depth analyses of *pol* sequencing data. A total of 373 overlapping genomic windows each of length 340 bp were selected, staggering the starting positions by 5 bp. For each window, a phylogeny was inferred for all read pairs that fully spanned that window, and ancestral state reconstruction divided the reads for each sample into distinct groups (subgraphs), with the *phyloscanner* Sankoff k parameter set to 12.5. A group of reads was flagged as likely contamination if it contained three or fewer reads, or less than 0.1% of the total number of reads from the sample in that window. The consensus sequence and minority base frequencies were then re-calculated from the resulting cleaned mapped reads using shiver [22]. The cleaned mapped reads were also reanalysed with *phyloscanner* for inference of transmission between sampled individuals [8, 24].

Finally, both the consensus sequence and the cleaned reads were analysed with the Stanford drug resistance tool [25] to determine consensus and minority drug resistance levels. Aggregated drug resistance predictions, accounting for mutations linked on the same read pair, were calculated as the maximum level of resistance (Susceptible < Potential low-level < Low-level < Intermediate < High-level) observed in at least 20% of merged read pairs spanning each position.

## 3. Results

### 3.1. Optimisation of existing methods

The starting point for this protocol was the established method, veSEQ, for genomic characterisation of Hepatitis C Virus [12]. Several issues limited the direct application of veSEQ to HIV. First, compared with HCV, HIV is more frequently found with viral loads below 10^4^ copies per ml, which is close to the threshold for reproducible whole-genome sequencing by veSEQ. Second, the veSEQ protocol was relatively laborious, with many steps and limited options for increasing throughput. Third, the veSEQ protocol involved fragmentation of the RNA in addition to the random priming of cDNA synthesis, resulting in short fragment size that would limit the use of the data for studies of within-host diversity. Minority variants are of particular importance for the identification of drug resistance and phylogenetically informed analysis of transmission [8]. Finally, all data generated by NGS suffer from issues of contamination, which need to be addressed for any application that analyses individual read sequences rather than the consensus of a large number of reads.

To address these limitations, we performed a series of optimisations (Supplementary Table 2), resulting in changes throughout the protocol. Briefly, our aims were to increase fragment size, maximise throughput, reduce sample processing time, simplify processing steps, and reduce reagent costs. Our final laboratory protocol consists of the following steps:

1. Viral particle lysis using chaotropic guanidine thiocyanate and total nucleic acid extraction using magnetised silica (easyMAG, bioMérieux).
2. RNA concentration and sample volume reduction using magnetised silica beads (RNAClean XP, Beckman Coulter).
3. Synthesis of libraries in low volume reactions, with low-temperature RNA denaturation and the Switching Mechanism At the 5’ end of RNA Template (SMART) technology [26] to convert RNA to double stranded Illumina libraries within a single tube reaction (Clontech).
4. Minimal PCR amplification and double-indexing of sequencing libraries using indexed primers to reduce risks of index miss-assigned reads and false minority variants being generated by template switching.
5. Pooling libraries by equal volume, rather than by equal mass, with reduced hands-on time.
6. Size selection of pooled libraries using a stringent cleanup with AMPure XP magnetised silica beads (dependent on the molecular-size distribution of the non-size-selected pooled sequencing libraries, assessed by gel electrophoresis)
7. Bait capture of virus sequences using a panel of oligonucleotide probes designed to capture the expected diversity of HIV in sub-Saharan Africa.
8. Parallel library production and sequencing of 384 libraries on a HiSeq Rapid instrument set to produce 250 nt paired-end sequences within a single batch.

The resulting raw data is processed using open-source computational tools to:

1. Provide quality control statistics to help the laboratory quickly detect and fix problems.
2. Remove unwanted information and contaminants from raw sequencing output files.
3. Infer a sequence-derived viral load.
4. Infer consensus genotypes, minority variants and minority haplotypes.
5. Infer transmission chains, with quantified statistical support for links and direction of transmission.
6. Infer drug resistance, both at the consensus and minority haplotype level.

### 3.2. Quantification of viral load

Viral load is a measure of the concentration of virus in a sample. It is usually measured with highly standardised and regulated clinical assays using quantitative PCR that amplifies both the material to be tested and spiked internal standards of known viral load. Consistent with our previous study of veSEQ applied to HCV, we found that in contrast to amplicon-based sequencing, veSEQ-HIV was quantitative, in that the amount of recovered sequence correlated with the viral load [27]. This arises because PCR conditions remain non-saturating and because of the unbiased nature of the probes used for virus enrichment. A further slight improvement is obtained by computationally removing duplicate copies of viral fragments from sequence data, which are generated by non-viral-specific PCR steps in the protocol. To assess quantitativity, we validated the protocol on a subset of 126 samples for which viral load measurements were obtained with a clinical assay. Figure 3A shows the correlation between the clinical viral load and number of viral fragments recovered during sequencing along with the R^2^ value (0.89). This correlation was robust over a wide range of viral loads (Figure 3B), including a subset that were below the quantifiable limit of the clinical assay (<50 copies per ml). The quantitative nature of veSEQ-HIV also implies that the inferred frequency of minority variants in sequence data is a good reflection of the corresponding frequencies in the sample.

**Figure 3:**
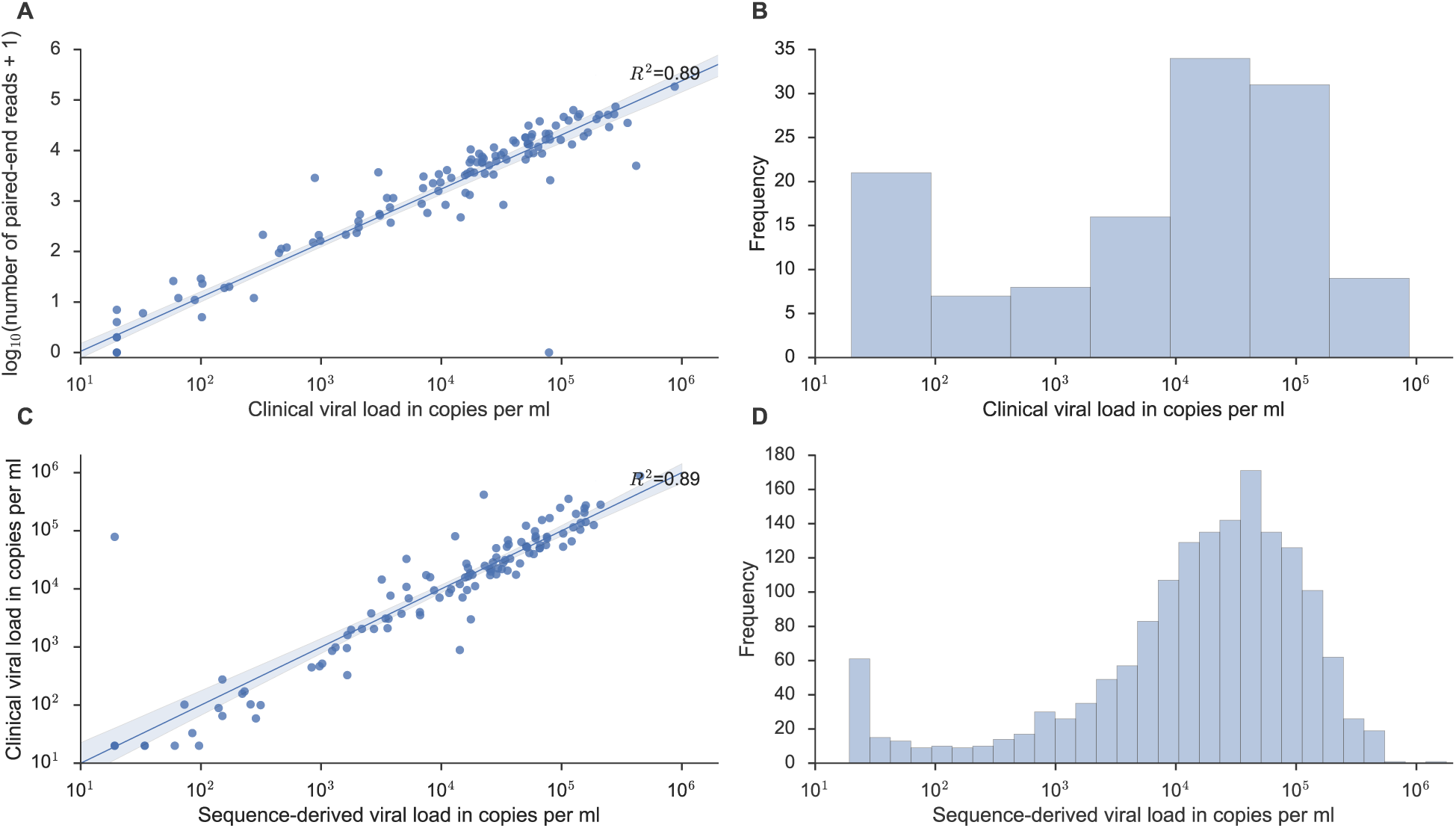
Panels A and B: Relationship between clinical viral load and number of mapped reads in a subset of 126 samples for which clinical viral load was measured independently; panels C shows the relationship between the clinical viral load and the sequence-derived viral load (using the equation in the main text) for these 126 samples. Panel D shows the frequency of sequence-derived viral load estimates for all 1,620 samples.

### 3.3. Relationship between clinical viral load and number of mapped reads

The relationship between number of reads and viral load was linear on a log-log scale with a slope of 0.83. This corresponds to some non-linearity on a linear scale, consistent with moderate bias of the sequencing method towards preferential recovery of the virus at low viral loads. This does not affect the use of the number of viral fragments to infer viral loads, since the relationship is well described mathematically.

Following the validation step, we defined “sequence-derived log viral load” as the linear transform of the log number of deduplicated sequence fragments (Figure 3C). The lower limit of detection is approximately 50 copies per ml. We calculated the sequence-derived viral load for all 1,620 sequenced samples using this transformation, and characterised the population distribution (Figure 3D). This distribution was bimodal, with the minor peak at very low viral load corresponding to individuals with HIV read counts below the quantifiable limit of conventional assays.

To ensure ongoing quantitativity and guard against batch effects, we introduced a panel of quantification standards, in line with the procedure used to calibrate clinical viral load assays. The standards comprised five dilutions of subtype B virus spiked into plasma (AcroMetrix HIV-1 Panel copies/ml, Thermo Fisher Scientific), and either one or two negative plasma controls. These were grouped with each batch of 90 HPTN-071-2 (PopART phylogenetics) samples at the point of RNA extraction. Although the sequence-derived viral load for this work was based solely on the clinically-quantified samples, the batch-specific standards can also be used to calculate the sequence-derived viral load and to quantify contamination within each sequencing run (Figure 4). We first introduced these standards in batch 6, and have been using these to monitor the quantitativity of each batch.

**Figure 4:**
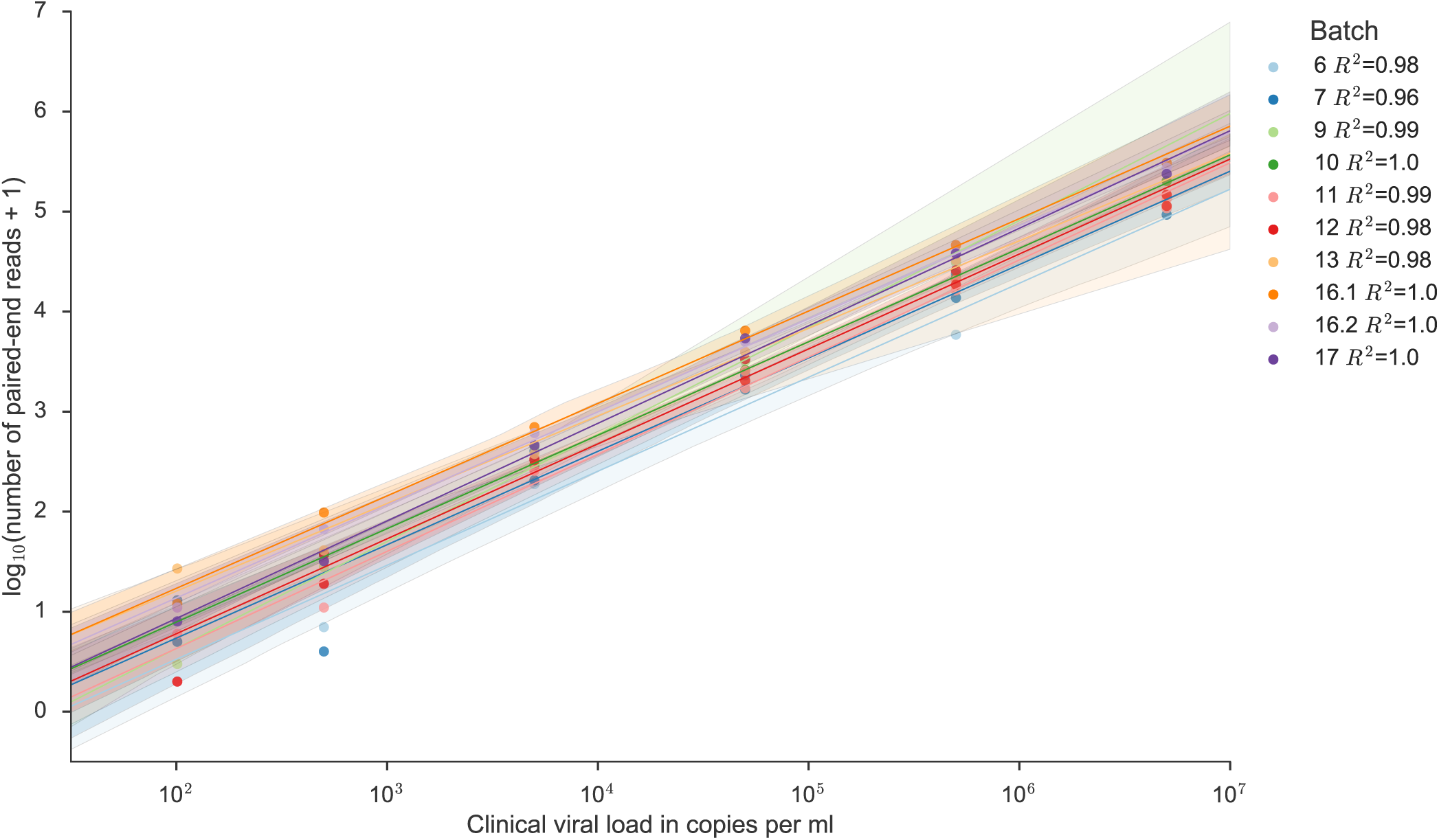
Within-batch quantification standards for calculating sequenced-derived viral load. Standard dilutions of subtype B virus were sequenced alongside HPTN-071 (PopART) samples, with the same dilution series in each batch. The figure shows the R^2^ values for linear regression lines fitted separately for each batch of standards. Shaded regions represent 95% confidence intervals with 1000 bootstrap replicates.

### 3.4. Genome coverage and sensitivity to viral load

The length of the consensus sequence, i.e. the amount of the genome that could be recovered from the reads, depends on the minimum read depth required to make a consensus call at each genomic position. Here, we define ‘read depth’ as the number of mapped reads covering each position in the genome after removal of PCR duplicates. To estimate the threshold for minimum read depth, we analysed our data with the tool LinkIdentityToCoverage.py in shiver, calculating the mean similarity between reads and the sample-specific consensus from shiver. The point at which reads consistently matched the sample consensus saturated at a depth of 5 reads, and we took this as our threshold for reliably inferring a consensus. This may be a conservative estimate given that sequencing negative controls resulted in no HIV reads after our multi-stage removal of contamination. However, we sought to produce not only accurate whole-genome consensus sequences, but also sufficient read depth for analyses of within-host diversity and characterisation of low-frequency drug resistance mutations.

Complete HIV sequences were obtained from the large majority of samples (Figure 5). The patterns of read depth were reproducible between individuals, with similar patterns of high and low coverage across the genome. Importantly, we did not observe a drop-off in coverage below 5 reads to be systematically associated with particular parts of the genome (Figure 6).

**Figure 5:**
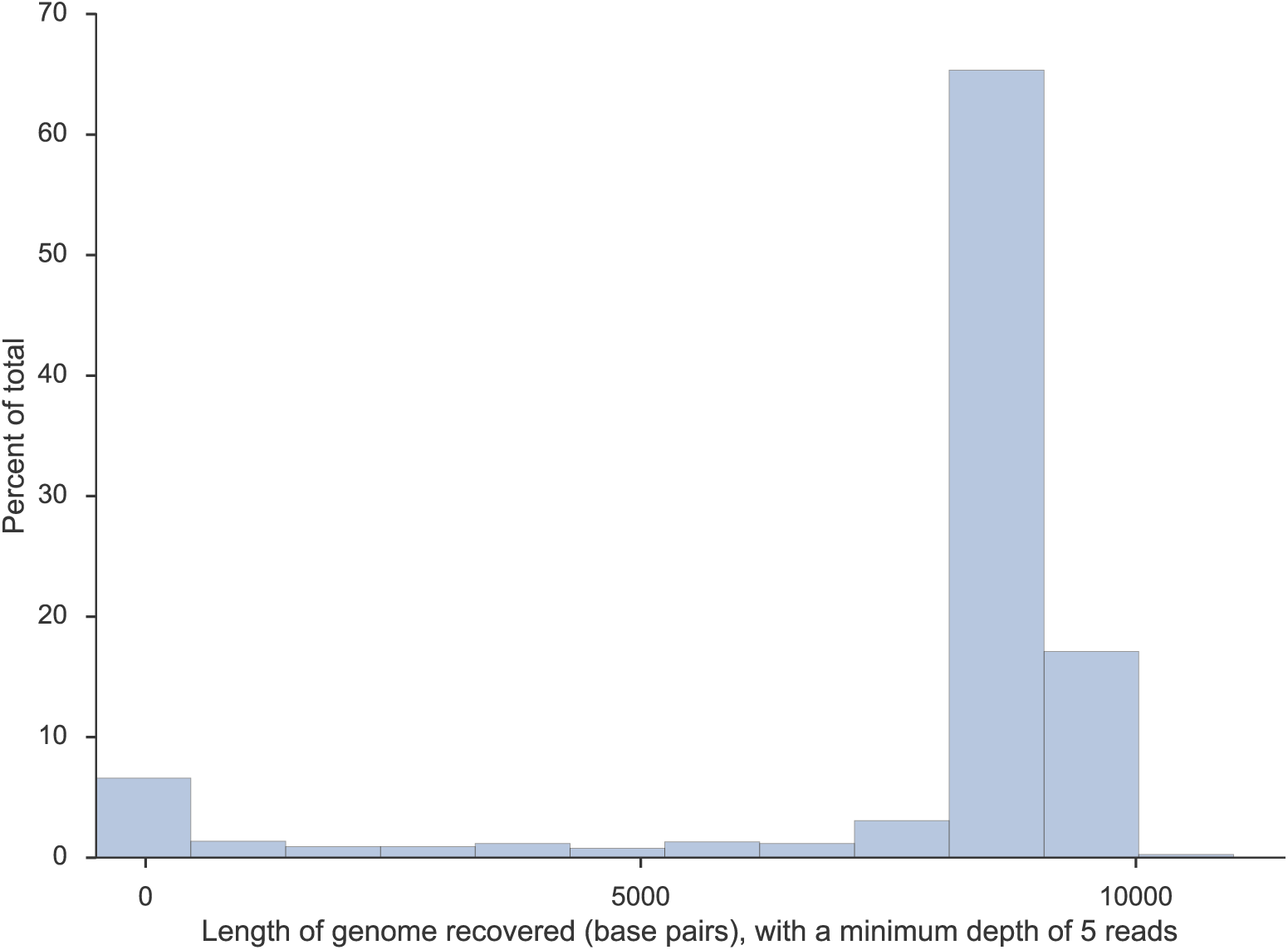
Length of recovered HIV genome for all sequenced samples. We consider a position in the genome accurately determined when the read depth is at least 5.

**Figure 6:**
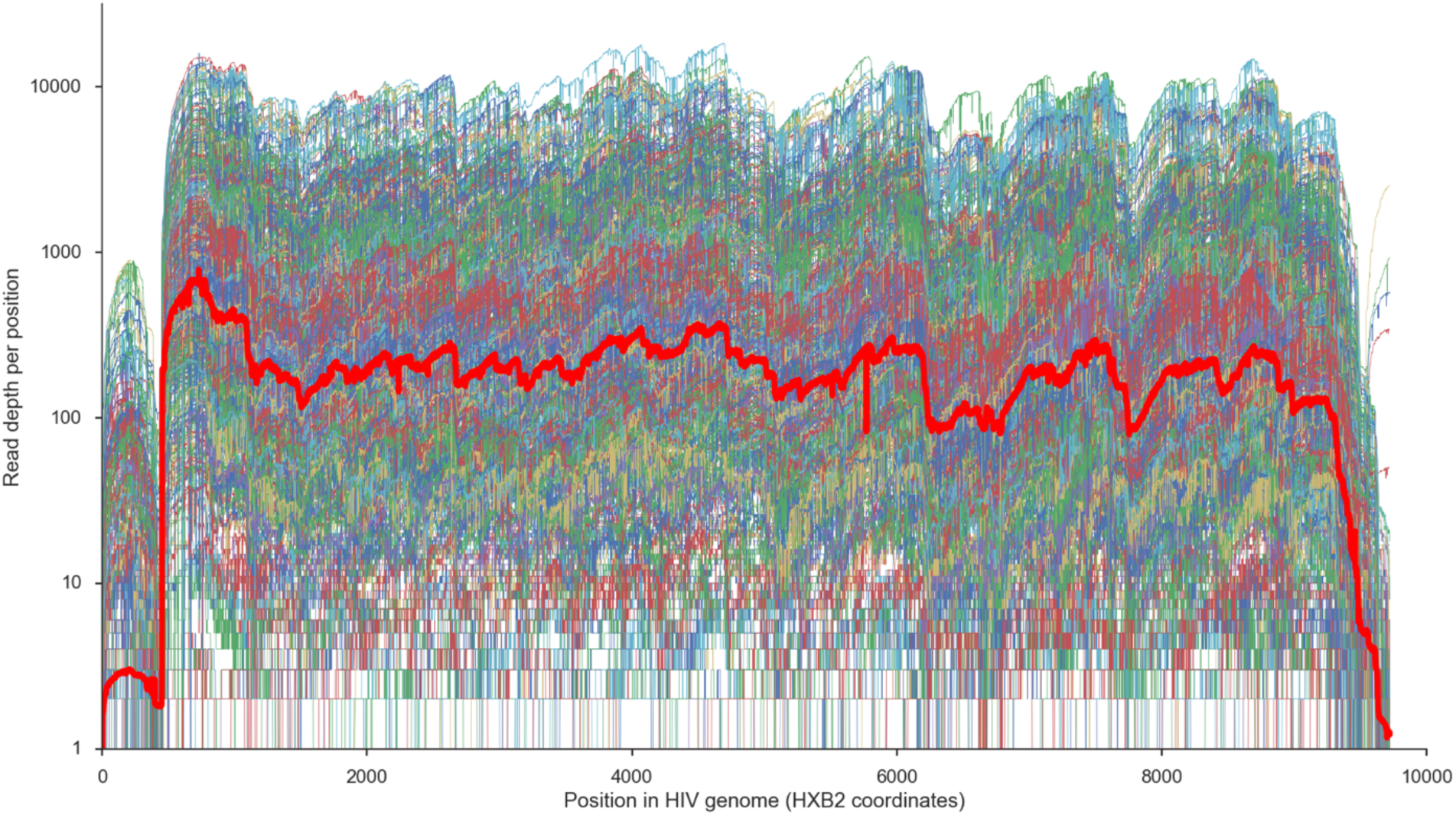
Read depth across the HIV genome for all samples in a single batch. Each coloured line corresponds to a single sample. The thick red line indicates the overall geometric mean for the batch. This is shown for the most recent batch (16_17_19, 456 samples), chosen for illustrative purposes.

Prior to batch 7, we used the original veSEQ approach for RNA fragmentation and cDNA generation followed by enzymatic-ligation of adapters for library preparation [12], which accounts for the different patterns of read depth observed in these batches (Figure 7).

**Figure 7:**
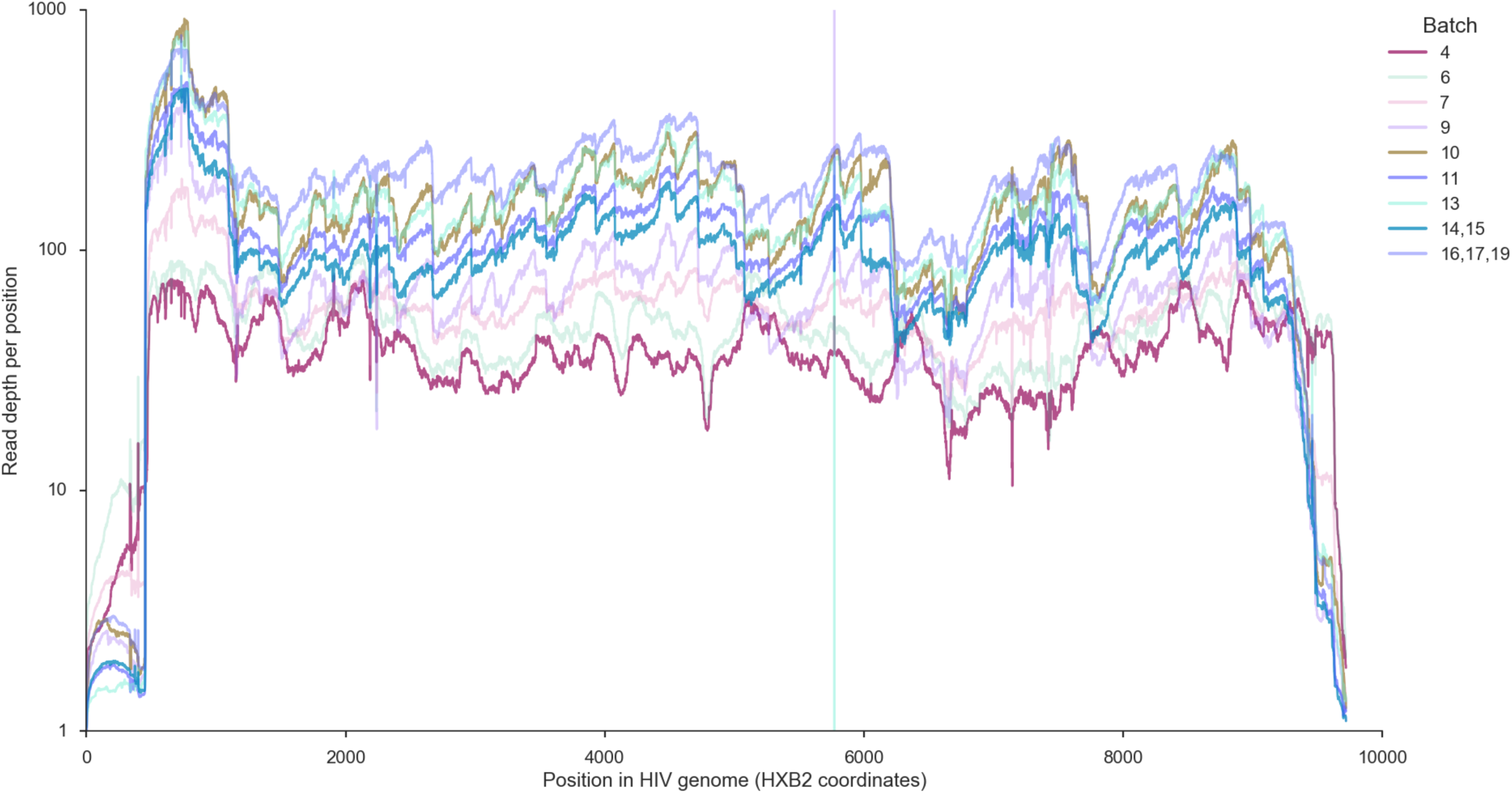
Mean read depth across the HIV genome for each batch of sequenced samples. Batches 4 – 6 were sequenced prior to the adoption of the full optimised protocol and did not include size selection of pooled libraries.

Following adoption of the SMARTer protocol from batch 7 onwards, we observed no significant batch effects. As discussed, we calculate viral load from the number of HIV reads obtained; Figure 8 shows that larger values of sequence-derived viral load (or equivalently larger total numbers of reads) are associated with a greater depth of reads across the whole of the HIV genome.

**Figure 8:**
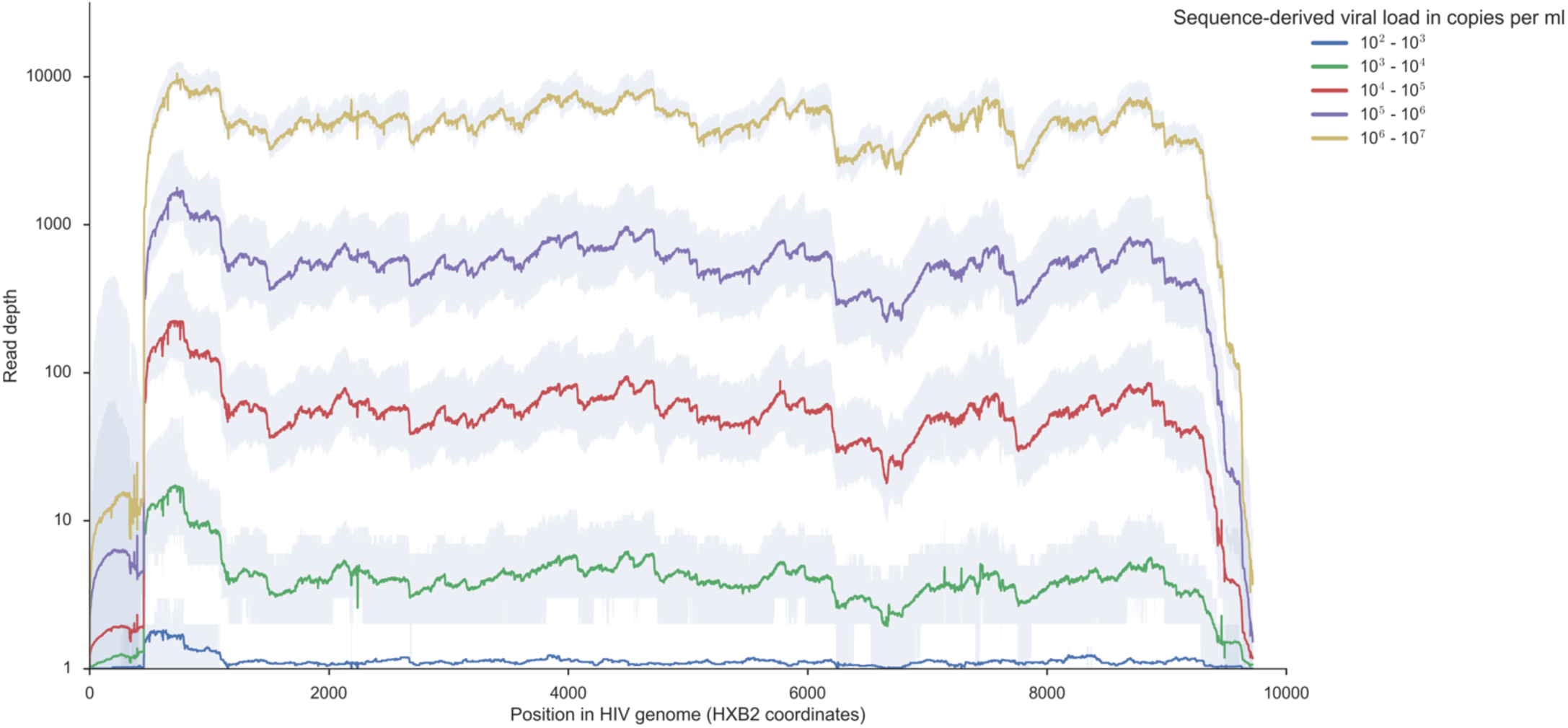
Mean read depth across the HIV genome by log sequence-derived viral load. Each line corresponds to all samples within the indicated log sequence-derived viral load range, with the shaded area showing the interquartile range. Higher read depth for samples with higher viral load results in greater coverage across the entire HIV genome.

Consequently, the success rate of reconstructing whole genomes increases with viral load; with the optimised veSEQ-HIV method we obtained complete genomes (defined as at least 8 kb) supported by a depth of at least 5 reads for 93% of samples with a viral load of 1000 copies per ml or more. Figure 9 shows the dependence of this success rate on viral load in more detail, for both the subset of samples with clinically validated viral load, and the full dataset with sequence-derived viral load. Sigmoid functions (fit to the data with least squares) indicate the viral load thresholds above which at least 8 kb genomes tend to be recovered: these are between 10^3^ and 10^4^ copies per ml, depending on the required depth of reads supporting the consensus.

**Figure 9:**
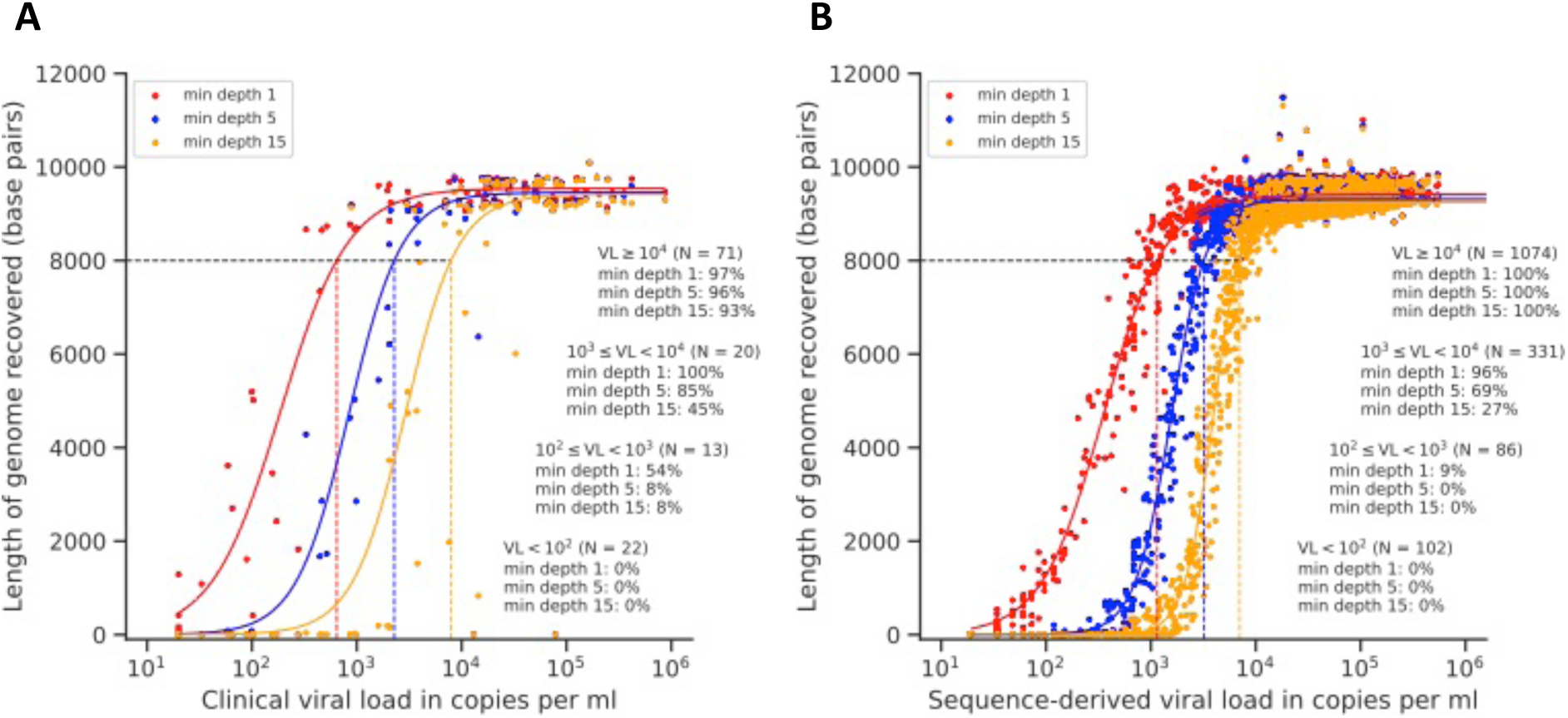
Genome recovery success rate as a function of viral load. Panel A, subset of 126 samples for which a clinically validated viral load measurement was available; Panel B, full data set using the sequence-derived viral load. For each sample we show three consensus genome lengths: those called requiring minimum read depths of 1, 5 and 15. The solid curved lines are least-squares sigmoid fits to the data. The dashed straight lines indicate the viral load values required to achieve a complete genome, defined as at least 8000 bp. The figures set within each plot report the fraction of samples with complete genomes, stratified by viral load.

### 3.5. Assay specificity

Complete HIV genomes, defined as sequence length over 8,000 bp with minimum read depth of 5, were recovered from HIV samples spanning the known HIV subtype diversity in Zambia (Figure 10). HIV subtypes were inferred by sequence similarity to HIV reference genomes from LANL; for a subset of genomes the subtype of the consensus was also inferred using the REGA HIV-1 subtyping tool [28]. The predominant subtype was C, for which 86% (1282/1498) of samples yielded complete genomes. Other subtypes of well-characterised complete genomes included A (A1 and A2), D, G, and J (Figure 10), as well as the subtype B standards, demonstrating good probe affinity across HIV diversity. Among possible recombinants (category “Other”), a lower proportion of samples yielded complete genomes 32% (6/19) — as expected given the difficulty in identifying the subtype of partial genomes. There was a strong relationship between the sequence-derived log viral load and genome recovery for all subtypes, consistent with viral load (and therefore total number of HIV reads)

**Figure 10:**
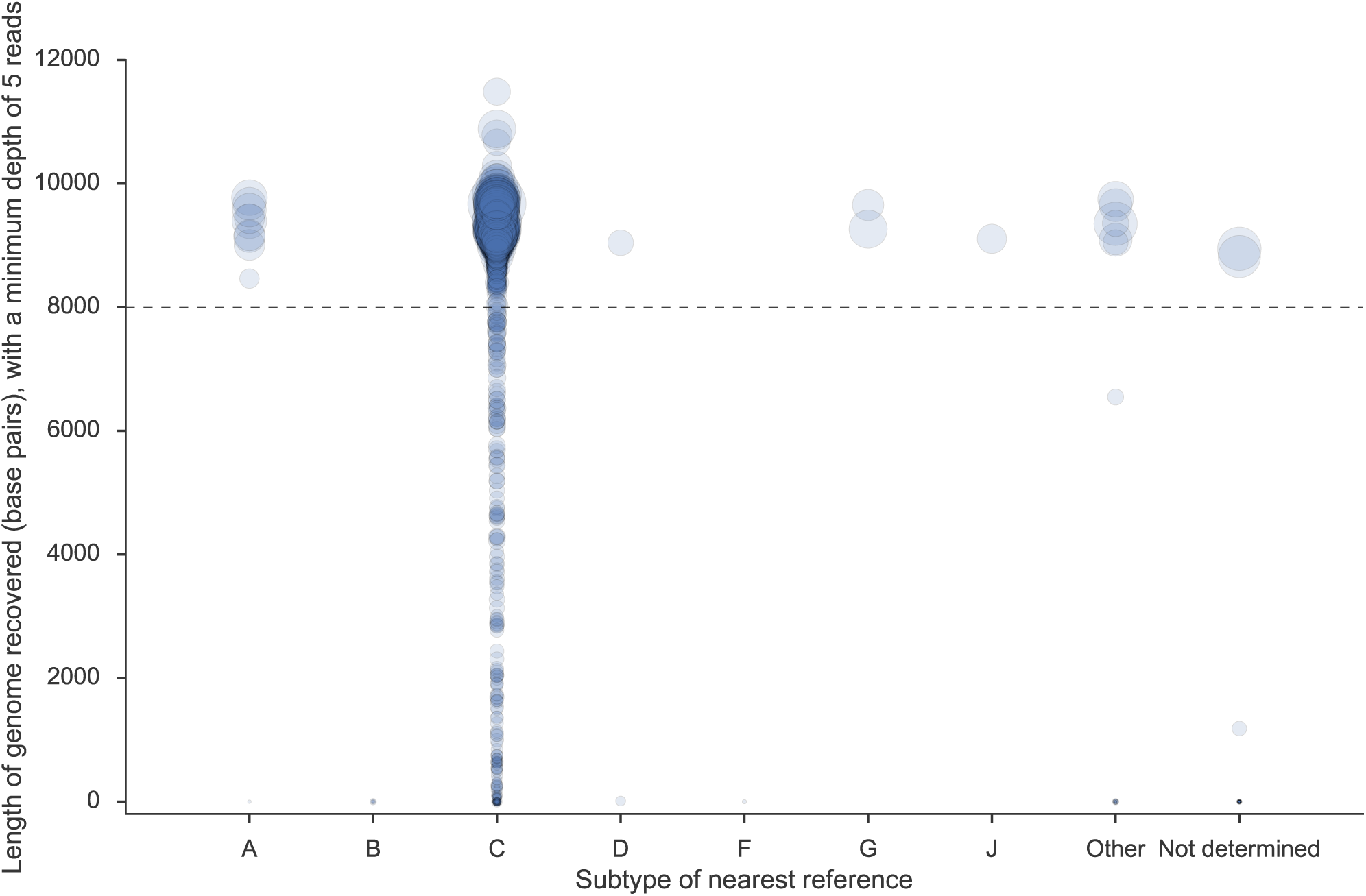
Probe sensitivity by HIV subtype and sequence-derived viral load. The category “Other” consists of potential inter-subtype recombinants. The marker size is proportional to sequence-derived log viral load. A minority of genomes (n=9, all subtype C) had a duplication of the repetitive LTR region in the consensus resulting in genome length greater than 10,000 bp; we have since modified our computational pipeline to prevent such artefacts. Quantitative standards (HXB2, subtype B) are not included in this analysis.

### 3.6. Contamination

Contamination can be physically introduced in the laboratory, or occur due to index misassignment errors during sequencing. It can undermine several important inferences, namely estimations of viral load (in particular distinguishing low viral load from aviraemia), the direction of transmission using within-host phylogenetics, and drug-resistant minor variants.

Phyloscanner contains several procedures not only for detecting contaminant reads in NGS datasets [8], but also for ‘blacklisting’ them (specifically removing them from consideration for further analysis). Blacklisting works by identifying reads in a sample that are either identical to those from a second sample but present in much smaller numbers, or by identifying reads that, while unique to a given sample, are relatively few in number and very phylogenetically distinct from the majority of the sample’s reads. For this analysis, we calculated the proportions of reads identified as probable contaminants in the 1,620 samples: Figure 11 shows the results. In panel A, each point represents one sample. Out of the total set of HIV reads obtained from all samples, 1.2% were identified as contamination; the mean percentage of reads per sample found to be contamination was 2.09%, reflecting variation in coverage between samples (for samples with few reads, small collections of reads which are suspected contaminants make up a larger proportion of the total). The complete workflow is included within *phyloscanner* (*phyloscanner clean*).

**Figure 11:**
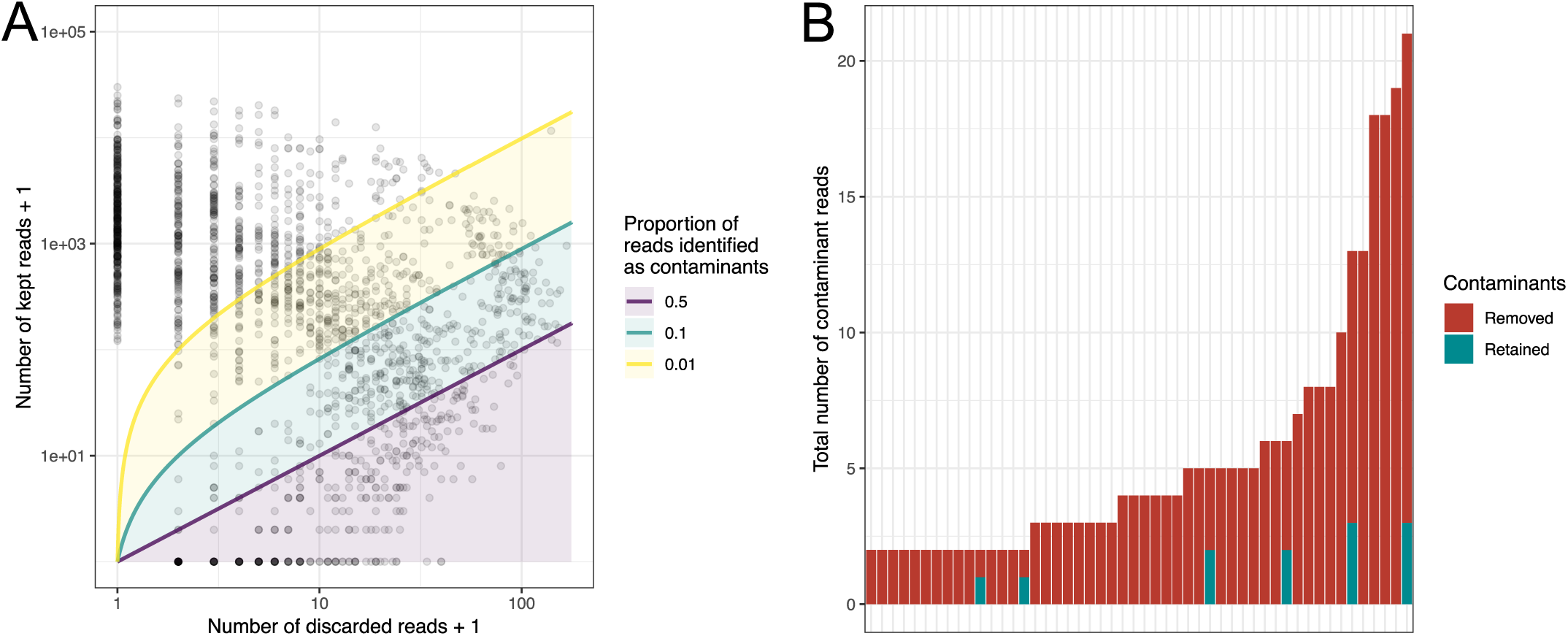
Panel A: Scatter plot of number of kept and blacklisted reads for 1,620 samples. A pseudocount of 1 has been added to values on both axes in order to allow use of a log scale. The coloured lines represent thresholds at which 50%, 10% and 1% of reads are identified as contaminants; samples appearing below each line have at least that many suspicious reads. The number of samples in each category were 343 (18.1%), 689 (36.4%) and 1088 (57.5%) respectively. Panel B: For 50 samples artificially contaminated by replacing 0.1% of their reads with the same number from another sample, the reads correctly identified as contaminants (red) and those not identified (blue).

We validated the blacklisting procedure on reads within the *pol* gene, which contains the majority of drug resistance mutations. We selected 100 samples with at least 2,000 reads in *pol* and cleaned each according to the method described. This set of 100 was divided into two groups of 50, and for each sample in the first group, 0.1% of reads were replaced with the same number of random reads from one member of the second group. We then tested phyloscanner’s removal of these known contaminant reads, using with the same settings as above. Cleaning removed 262 out of 274 contaminant reads, giving an overall sensitivity of 95.6%; the distribution of these reads over the 50 samples is shown in Figure 6B. 73 of 291,815 non-contaminant reads were identified as contaminants, giving an overall specificity of over 99.9%. Knowledge of the actual proportion of contaminant reads in a simulation exercise of this sort allows phyloscanner settings to be tuned for optimal performance.

### 3.7. Insert size

veSEQ-HIV aims not only to accurately determine the consensus genome that is, the most common base in the viral quasispecies at each position in the genome but also to accurately determine the variability between viruses in the quasispecies. Like most high-throughput NGS protocols, veSEQ-HIV requires fragmentation of the virus RNA into so-called ‘inserts’; we sought to optimise the length of these inserts to be as long as possible within the limits of what the sequencing machine was able to process. In previous work we found that inserts of 350 bp or more offer useful insights into within-host phylogenetic diversity [8]. Figure 12 shows how veSEQ-HIV was optimised to consistently generate over 40% of inserts above this desirable length.

**Figure 12:**
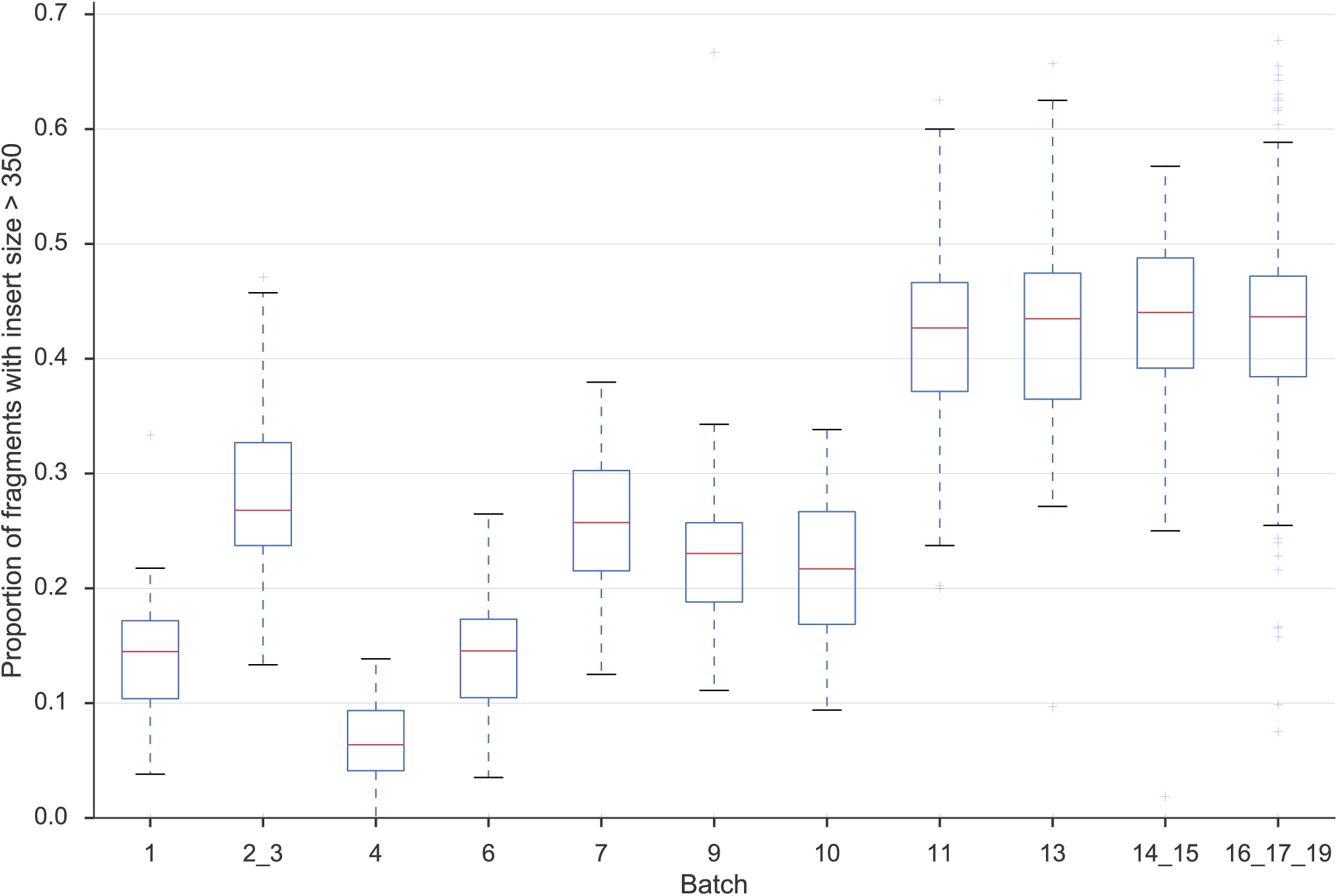
Optimisation of the proportion of sequenced fragments longer than 350 bp. Batches are presented in chronological order, spanning a period of 18 months. The SMARTer protocol was introduced with batch 7; bead-based size selection was introduced at batch 11; reagent volumes were scaled down for batch 16_17_19 with no detrimental effect on the proportion of all deduplicated fragments greater than 350 bp in length.

### 3.8. Cost

The cost of implementing a high-throughput virus genomics system will vary by setting, depending on many local factors. In our laboratory in Oxford, the reagent and consumables cost of the entire assay, from frozen blood to final data, is approximately 45 USD in 2018, 50% lower than the cost of a locally obtained clinical viral load.

### 3.9. Drug Resistance

The observation that sustained pressure from suboptimal treatment with ART invariably selects multiple mutations at the consensus level is evidence that linkage between mutations increases the overall replicative fitness of a virus exposed to treatment. If low-frequency mutations are observed at multiple different positions, Sanger consensus sequencing does not tell us whether they occur together (we know only that there are some minor variants at one position and some at the other, but not whether these are found within the same viruses). Hence, we cannot distinguish whether there is a variety of different viruses each with a low level of drug resistance due to a different mutation, or a small subset of viruses with multiple resistance mutations, giving a higher overall level of resistance. Making this distinction may be relevant to predictions of treatment success and the fixation and transmission of mutations within populations. Furthermore, because the consensus sequence represents average information at each base, minority nucleotide substitutions occurring within the same codon can lead to spurious predictions of the true amino acids.

NGS resolves these problems by providing information on linkage between mutations located on the same sequenced fragment. Figure 13 shows an example of the genetic and phenotypic diversity of reads spanning regions of the *pol* gene that contain several sites affecting resistance to the NNRTI and NRTI classes of ART drugs. The sample from patient A contains reads with as many as 6 mutations co-occurring on the same sequenced molecule. We calculate the overall phenotype for an individual as the maximum level of drug resistance in reads across the *pol* gene. We require any given mutation to be present in at least 20% of clean reads, after removal of contamination and PCR duplicates as described. In the same individual, the most susceptible virus appears closest to the root of the phylogenetic tree (i.e. is closest to the founder virus), which suggests accumulation of resistance mutations during the course of infection. Consistent with this, the individual took ART prior to sample donation.

**Figure 13:**
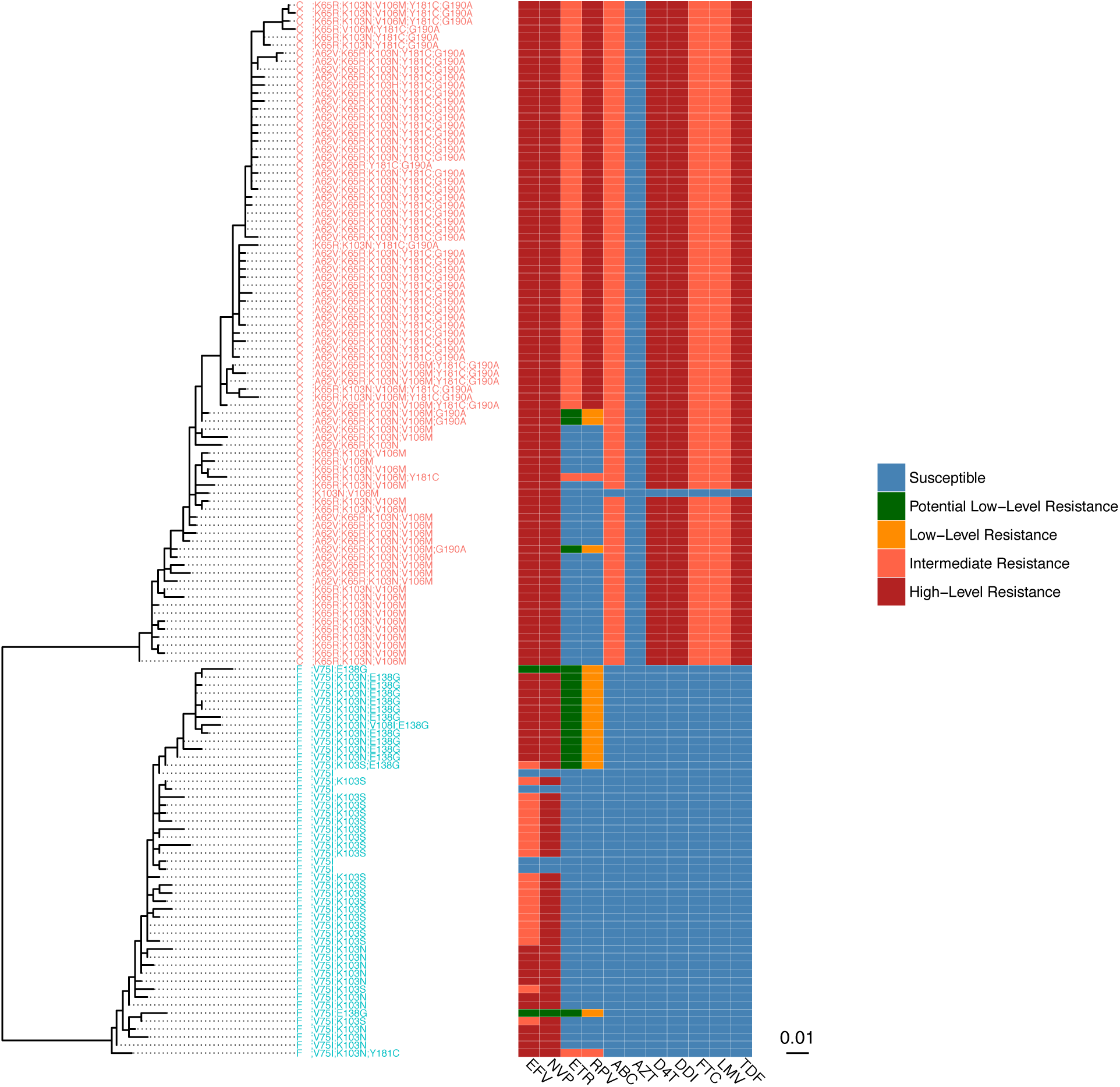
Within-host diversity in two patients with differing patterns of drug resistance. The phylogenetic relationship (left) and drug resistance results (right) inferred from 350 bp alignments of the polymerase gene for two patients. Each tip in the phylogeny and line in the resistance plot corresponds to a unique viral genotype recovered from the patient’s quasispecies. The predicted level of resistance to drugs from the NRTI and NNRTI classes (bottom) are shown.

### 3.10. Discussion

We have developed, optimised and validated veSEQ-HIV, a complete high-throughput laboratory and computational process for recovering complete HIV genomes including minority variant frequencies, estimating viral load, and detecting ART drug resistance (Figure 14). The method has been shown to work with 1,620 genetically diverse samples collected from 10 Zambian clinics participating in the HPTN-071 (PopART) trial, producing complete genomes from >90% of samples with viral loads >1,000 copies per ml. The assay works with residual plasma taken from routine CD4 count testing obtained in field conditions, without introducing undue contamination or degradation of the samples or the need for additional blood draws.

**Figure 14:**
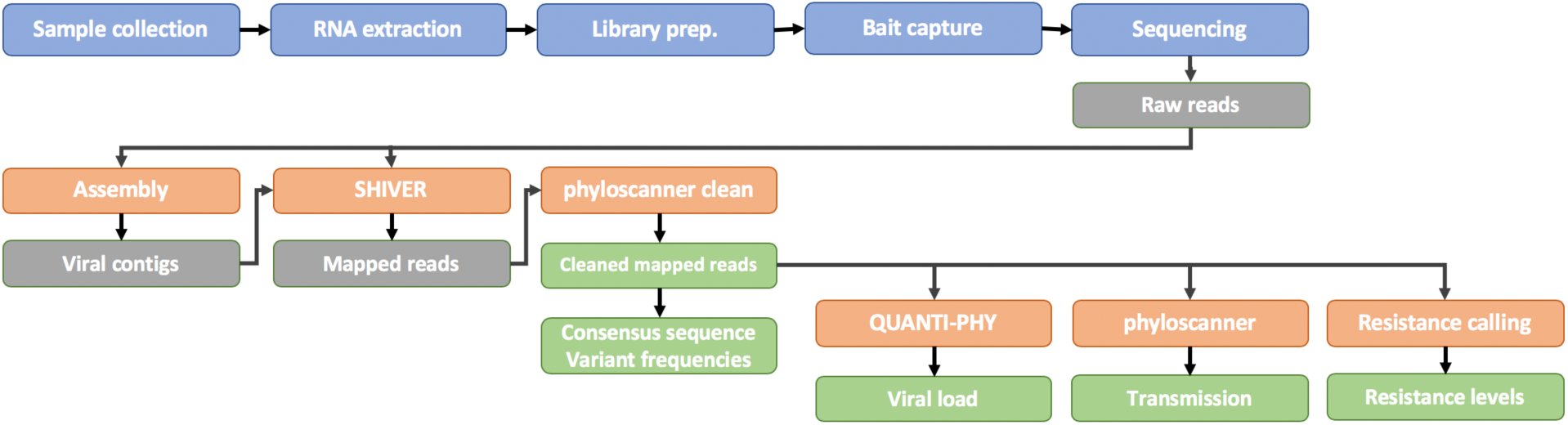
Overview of veSEQ-HIV: a complete laboratory and computational pipeline for high-throughput sequencing. RNA extraction from plasma samples is carried out in a CL-3 certified laboratory, before transfer to a dedicated genomics facility for library preparation, bait capture, and finally sequencing. Raw sequencing data is pre-processed to remove host and contaminant RNA, and these computationally filtered reads togetherwiththeir *de novo* assembled contigs are used to determinethe consensus genome and minority variant frequencies using shiver. QC metrics are then calculated, and the proportion of contaminant reads originating from other samples is estimated with Kallisto. Samples which result in a successful read mapping are then cleaned with phyloscanner with OptiPhy settings to remove contaminant reads, and clean reads are used to infer transmission patterns with phyloscanner, and to make drug resistance predictions with HIVdb.

Sequence-derived viral load estimated by veSEQ-HIV is cheaper than some existing commercial viral load tests, and is highly reproducible. Reported R^2^ values between commonly used clinical viral load assays range from 0.80 to 0.94 [29]. The veSEQ-HIV assay compared with the TaqMan assay, validated here on 126 samples, which gave an R-sq value of 0.89, well within this range. The detection and quantification limits of veSEQ-HIV (50 – 100 copies per ml) are likewise comparable with those of clinical viral load assays. Although the sample size for this validation exercise is small, we show that quantification is reproducible between runs by inclusion of quantification reference standards derived from an HXB2 isolate, diluted in normal human plasma. In future work, more clinical viral loads will be cross-referenced across several of sequences runs in order to confirm that a single clade B HXB2 isolate is a suitable quantification standard for other (mostly clade C) isolates. We expect this to be the case because we have previously shown bait capture performs without significant bias across a range of HCV subtypes, greater than the diversity of HIV observed here [12].

The throughput of veSEQ-HIV is suitable for large-scale public health applications. In our research setting a single technician is able to process 360 samples per week, and with use of a 384-well low-volume liquid handler, we can prepare the same number of samples in a day. Routine combination testing to provide vital information on viral suppression, drug resistance and transmission, in near real time, is entirely possible with veSEQ-HIV.

The aim of our ongoing work is to provide insights into the outcome of the HPTN 071 (PopART) trial [15]. Analysis of viral genetics will be used to estimate the proportion of transmissions that occur during early and acute HIV infection, the proportion of transmission events that occur from individuals living within or outside of the trial communities, and the demographic and epidemiological correlates of being a transmitter. All analyses will take place in both intervention and control communities. Similar analyses are planned for other recent trials of treatment as prevention [30]. Combining these data, for example in the PANGEA consortium [31] [http://pangea-hiv.org], will provide further insights into the generalisability of findings, and larger scale patterns of viral flow and human migration.

As well as being cost-effective, our method has several advantages over previous high-throughput approaches [9]. Because our method minimises the biases involved in PCR, and computationally controls contamination, our estimates of the frequency of minority genetic variants are likely to be more robust. Second, the design of our probes is unbiased with respect to the range of commonly found HIV viral variants. Abbott Laboratories has recently reported on a similar method, developed in parallel to ours, which they show works across a wider panel of reference genomes [32, 33].

There remain important limitations to our approach. Whilst we have validated our approach to minimise contamination, and provide QC tools to detect mix-ups as quickly as possible, the risk of large-scale mix-ups increases with higher throughput. This should be mitigated with sample barcoding and sample tracking. Second, our results remain to be independently clinically validated: veSEQ-HIV is not a licensed viral load, genotyping or drug resistance assay. Cross validation against licensed drug resistance assays is a priority.

### 3.11. Future directions

The veSEQ-HIV protocol is tuned for high-throughput applications, and so is ideally suited for laboratories that process a large number of samples. Capital investments are modest, and the protocol is simple for technicians to adopt. Maintenance and supply issues could be problematic in lowand middle-income countries, where the need is greatest.

In separate experiments, we successfully sequenced bait-captured viral genomes on the portable Oxford Nanopore instrument in a single day, both in our laboratory and in a semimobile BSL3 laboratory in Zambia. Read lengths are longer; however, as has been reported elsewhere, the error rate in single reads is high. We are pursuing strategies to correct error by repeatedly sequencing linked fragments derived from single templates, but presently, due to limits to multiplexing, costs per sample for Nanopore are substantially higher.

Future areas for improvement might include increasing automation, reducing initial capital expenditure costs, and reducing the reliance on regular supply chains of consumables. We did not explore the extent to which the bait capture step could be shortened or simplified; such improvements would further simplify the implementation of our method and would be needed to achieve a high-throughput protocol that could turn around sequence data in a single working day. Extending the length of individual sequences to capture whole viral haplotypes would improve applications in epidemiology and pathogenesis research.

The method can easily be adapted to study other RNA viruses or a panel of RNA viruses without loss of sensitivity. Since oligonucleotide probes are typically a one-off investment, sequencing several pathogens does not substantially increase the cost of the assay.

The computational component of the method is currently optimised for our local cluster infrastructure, and needs to be streamlined and made platform-independent. It is also possible to tune performance and port the code to be run either on secure cloud services or on standalone machines with reduced computational burden. Patient groups should be regularly consulted on the ethical use of this technology, providing maximal benefits whilst minimising the risks (Coltart et al., Lancet HIV in press).

In summary, we have reported a cost-saving high-throughput protocol that, with current technologies produces a sequence-derived viral load, a high-resolution drug resistance genotype, and data that can be used to provide highly granular insights into HIV epidemiology. The method has proven robust to field conditions in Zambia and carries no additional testing burden for patients: we used residual blood from routine CD4 tests. Sequencing will provide insights into the outcome of the HPTN 071 trial, and in our view, could be included with much higher coverage than is commonly done in future epidemiological and intervention studies of RNA virus spread.

## Acknowledgements

Sponsored by the National Institute of Allergy and Infectious Diseases (NIAID) under Cooperative Agreements # UM1 AI068619, UM1-AI068617, and UM1-AI068613. Funded by: The U.S. President’s Emergency Plan for AIDS Relief (PEPFAR), The International Initiative for Impact Evaluation (with support from the Bill & Melinda Gates Foundation), NIAID, the National Institute of Mental Health (NIMH), the National Institute on Drug Abuse (NIDA). We thank Monique Andersson and the John Radcliffe Hospital, Oxford, Clinical Microbiology Department for assistance with viral load testing. We acknowledege the support of the HPTN 071 (PopART) study team and Zambian Ministry for Health. Sequencing was supported by the Oxford Viromics initiative (Paul Klenerman, Rory Bowden) and the Oxford Genomics Centre (With thanks to John Broxholme, Lorne Lonie, Angie Green and David Buck). Sample and data collection has been supported by the PANGEA HIV consortium funded by the Bill & Melinda Gates Foundation.

